# Inferring rates of metastatic dissemination using stochastic network models

**DOI:** 10.1101/352096

**Authors:** P. Gerlee, M. Johansson

**Affiliations:** Mathematical Sciences, Chalmers University of Technology and University of Gothenburg, Sweden; Department of Oncology, Institute of Clinical Sciences, Sahlgrenska Academy at the University of Gothenburg, Sahlgrenska University Hospital, Gothenburg, Sweden

## Abstract

The formation of metastases is driven by the ability of cancer cells to disseminate from the site of the primary tumour to target organs. The process of dissemination is constrained by anatomical features such as the flow of blood and lymph in the circulatory system. We exploit this fact in a stochastic network model of metastasis formation, in which only anatomically feasible routes of dissemination are considered. By fitting this model to two different clinical datasets (tongue & ovarian cancer) we show that incidence data can be modelled using a small number of biologically meaningful parameters. The fitted models reveal site specific relative rates of dissemination and also allow for patient-specific predictions of metastatic involvement based on primary tumour location and stage. Applied to other data sets this type of model could yield insight about seed-soil effects, and could also be used in a clinical setting to provide personalised predictions about the extent of metastatic spread.

**AUTHOR SUMMARY:** For most cancer patients the occurrence of metastases equals incurable disease. Despite this fact our quantitative knowledge about the process of metastatic dissemination is limited. In this manuscript we improve on a previously published mathematical model by incorporating known biological facts about metastatic spread and also consider the temporal dimension of dissemination. The model is fit to two different cancer types with very different patterns of spread, which highlights the versatility of our framework. Properly parametrised this type of model can be used for making personalised predictions about metastatic burden.

## INTRODUCTION

For many forms of cancer, occurrence of distant metastases equals incurable disease [1, 2]. Regional metastases, i.e. positive lymph nodes, also implies an inferior prognosis [3, 4]. In order to diminish the risk of dissemination of the primary tumor, patients commonly receive adjuvant treatment with radiotherapy and/or some kind of medical oncological treatment. Yet, in many cases, the patient is later on faced with residual or recurrent disease [5]. On the other hand, many patients receive adjuvant treatment with subsequent side effects, even though their illness would never have disseminated [6, 7]. Hence, increased knowledge about the extent of metastasis at diagnosis would improve the care of many cancer patients.

Metastatic spread is known to follow certain disease specific patterns [8, 9], but despite the current state of knowledge, a quantitative understanding of the process of metastatic spread would improve our ability to optimise therapy and reduce over-treatment. Here will we make use of stochastic modelling to quantitatively assess the importance of different routes of dissemination. In addition, this method also makes it possible to estimate the extent of metastatic spread based on tumour stage and location. Our approach is versatile and we highlight this by applying it to two different types of tumours: carcinoma of the oral tongue, which disseminates primarily through the lymphatic system and ovarian carcinoma, which spreads both intraperitoneally, lymphatically and through the blood circulatory system.

The tendency of a primary tumour to form a metastasis is the hallmark of malignant cancer [10], and the process by which this occurs is a multi-step process [11, 12]. Typically tumour cells detach from the primary tumour, invade the surrounding stroma and find their way to local lymph nodes or blood vessels [13]. Although a late stage tumour can release very large number of circulating tumour cells (CTCs) into the blood stream (up to 4 × 10^6^ cells shed per gram of tumour per day [14]), the low probability of forming metastatic foci [15] coupled with the low probability of passing through capillary beds [16] lead to the conclusion that micrometastases in for example the liver (for gut malignancies) and the lung (for all malignancies) are necessary for further hematogenic dissemination [17–19]. These, often microscopic [20], lesions release CTCs into the arterial side of the circulation (in the case of the lung)[21], and hence amplify CTC numbers in arterial blood, which otherwise would be low due to the filtration occurring in the lung capillary bed. A similar process is at work during lymphatic spread [22], where CTCs get trapped in lymph nodes, where they form micrometastases, which shed CTCs that travel further downstream in the lymphatic system.

This process is known as secondary seeding, and although it is of paramount importance for the metasatic process we still have a limited understanding of the steps involved. Here mathematical modelling can provide a helping hand, since it allows for a quantitative understanding of biological processes that are difficult to measure directly.

Mathematical modelling of metastasis dates back to a series of seminal papers by Liotta and coworkers [23–25]. They considered the release of CTCs from an implanted tumour in mice and the subsequent formation of lung metastases. Using both deterministic and probabilistic methods they could derive predictions of how the number of metastatic foci changes over time and how the probability of being free of metastasis changed [24]. These predictions agreed well with experimental data and highlights the stochastic yet predictable nature of metastatic spread.

Another important contribution was made by Iwata et al. [26] who formulated and analysed a model which accounts for secondary seeding. That model predicts how the size distribution of metastases changes as the disease progresses. Predictions of the model were tested by Baratchart et al. [27] in a murine model of renal carcinoma, and while the model could predict the total metastatic burden, it was unable to describe the size distribution of metastases. However, the model prediction could be improved by assuming interactions between metastatic foci.

The Iwata model has also been used for connecting presurgical primary tumor volume and postsurgical metastatic burden and survival [28]. The model was applied to two datasets from mouse models and one clinical dataset and the analysis revealed a highly nonlinear relationship between resected primary tumor size and metastatic recurrence.

A similar approach has been used by Hanin and coworkers in a model which accounts for the growth rate of the primary tumour, shedding of metastases, their selection, latency and growth in a given secondary site [29]. They proved that the parameters of the model are identifiable in the case of Gompertzian growth of the primary tumour, and apply a maximum likelihood method to identify the model parameters in the case of a single patients with a large number of lung metastases.

Other mathematical models have successfully been applied in order to describe the dynamics of metastatic spread in pancreatic cancer [30] and breast cancer [31].

The importance of secondary seeding was investigated by Scott et al. [19] in the context of self-seeding [32], the process whereby a primary tumour can accelerate its own growth by releasing CTCs that return to the site of origin. A careful mathematical treatment of this hypothesis showed that secondary seeding is indeed required for this pathway to contribute to primary growth.

Secondary seeding also has an impact on estimates of metastatic efficiency, since undetected micrometastatic lesions render the apparent spread directly from the primary to target sites more efficient than it actually is [18].

The idea of metastatic spread occurring on a network, where the nodes represent organs and the links correspond to routes of spread was first described by Scott et al. [33], and later modelled quantitatively by Newton et al. [34, 35]. They considered a stochastic model where the dissemination of cancer cells is modeled as an ensemble of random walkers on the network. The dynamics of the model is determined by a transition matrix, which was obtained by fitting the model to a large autopsy data set [36]. The entries of this transition matrix gives information about rates of dissemination between different organs.

Here we build on the work of Newton et al., but with secondary seeding in mind, make modifications which alleviate the problem underdetermination, which plagued their work. In order to parametrise their model they had to assume that the observed patterns of metastasis correspond to a steady-state distribution of the model. We instead make use of primary tumour stage to create a model which is temporal and considerably easier to parametrise.

Focusing on two primary tumours we show that our model is able to estimate the dissem ination rates with high confidence, and crucially allows us to estimate rates between sites, which are inaccessible if one simply analyses incidence data.

## RESULTS

Our aim is to quantify the rate of metastatic spreading by applying a stochastic network model to clinical data. We will consider two different data sets: (i) A cohort of 141 patients diagnosed with tongue cancer where metastases occur in the head and neck region and (ii) data from patients with ovarian cancer obtained from the SEER-database where metastases occur both regionally and to some extent in distant organs. Regional spread typically occurs via the lymphatic system and metastases appear in lymph nodes (LN), whereas distant metastases are mediated by the blood circulatory system and appear in other organs such as the liver or the lung. Although the two processes are different in some respects they also share many commonalities. Firstly, the spread of the disseminated tumour cells is constrained by anatomical structures, i.e. the lymphatic system and the blood circulatory system. Secondly, the spread is directed since it is subject to the flow in the system, and lastly, the formation of metastases at secondary sites affects the downstream rate of formation since metastases, like the primary tumour, disseminate tumour cells as they grow. These similarities makes it possible to formulate a general mathematical model, which can be tailored to describe both data sets.

The model consists of a network with sites and directed links [33]. The nodes correspond to lymph node stations or organs, and be in either of two states: negative (i.e. containing no metastases) or positive (containing one or more metastases). The state of each node is a random variable, and the probability of it being positive depends on the state of the other nodes, and the rates of shedding between the nodes.

We assume that a positive site sheds CTCs that flow according to the links of the network. This applies both to the primary tumour and subsequent metastases. For simplicity we do not consider the size of metastases in this model, since this would require more parameters [37], and would make it more difficult to fit the model to clinical data. This implies that the state of each site is a binary variable taking the value 0 if the site is empty and 1 if it contains a metastasis.

We assume that each cancer cell which is disseminated from a site has the same probability of forming a metastasis at a downstream site. Since it is known that filtration rates are high (only 1 in 10 000 cells pass through a capillary bed) [19] we assume that CTCs only flow to neighbouring sites, e.g. CTCs released on the venous side of the circulatory system can only give rise to metastases in the lung, since this is the first capillary bed they encounter. This is also a good approximation for lymphatic spread, where the occurrence of skip metastases, in which intermediary LNs are negative, is rare [38], and has been suggested to be the results of undetectable micrometastases [38]. We aggregate the rate of release of CTCs, the survival probability in the circulatory system and the probability of forming a metastasis in a downstream site into a single rate parameter, which we call λ, when flow is from the primary tumour, and ϕ, when flow is from metastatic sites. This parameter then corresponds to the rate at which a downstream site becomes positive given a positive site upstream. We assume that the primary tumour (and secondary lesions) start shedding CTCs upon formation, which means that we assume a parallel progression model [39], as opposed to a linear model where shedding from the primary is delayed and occurs after malignant progression.

We include a temporal dimension into the model by using the primary tumour stage as a proxy for time. This means that we consider the stage as informative about the total number of CTCs that have been disseminated from the initiation of the tumour up until diagnosis. This is of course a crude approximation since it is known that tumours progress at different rates and preferentially disseminate to different sites depending on for example which mutations they harbour [40]. Although tumours might progress at different rates they still have to pass all the intermediary stages, and very rarely regress to a lower stage. If we knew the time since tumour initiation for each patient we would able to infer the rates in absolute terms (e.g. units per year). However, since we only have information about the tumour stage, and in addition do not have a mapping from real time to tumour stage, the rates that we infer from clinical data are only relative, but given a primary tumour of a certain stage the model can still predict the probability of metastases at different sites. The relative magnitude of the rates also informs us about the risk of developing new metastases given a certain metastatic burden. This approach was also taken by Benson et al. who used stage as a proxy for time in a model of metastatic spread in head and neck cancers [41]. In order to account for the fact that tumour volume, and therefore stage, typically changes non-linearly with time we also infer the flow rates under the assumption of exponential tumour growth (see Methods).

We now move on to discuss the models specific to tongue cancer and ovarian cancer.

### Lymphatic spread of tongue cancer

The drainage of the lymphatic system in the head and neck area can be described by a network where the nodes correspond to lymph node stations and the links represent flow between the stations. The location of the primary tumour determines how much it drains into the different stations, Tumours of the tongue mainly drain into station I and II, and to some extent to station III. Station I drains into II, which in turn drains into III, which drains into IV [38]. The dissemination network is shown in fig. 1, where the directed links show how CTCs flow in the system.

**FIG. 1.**
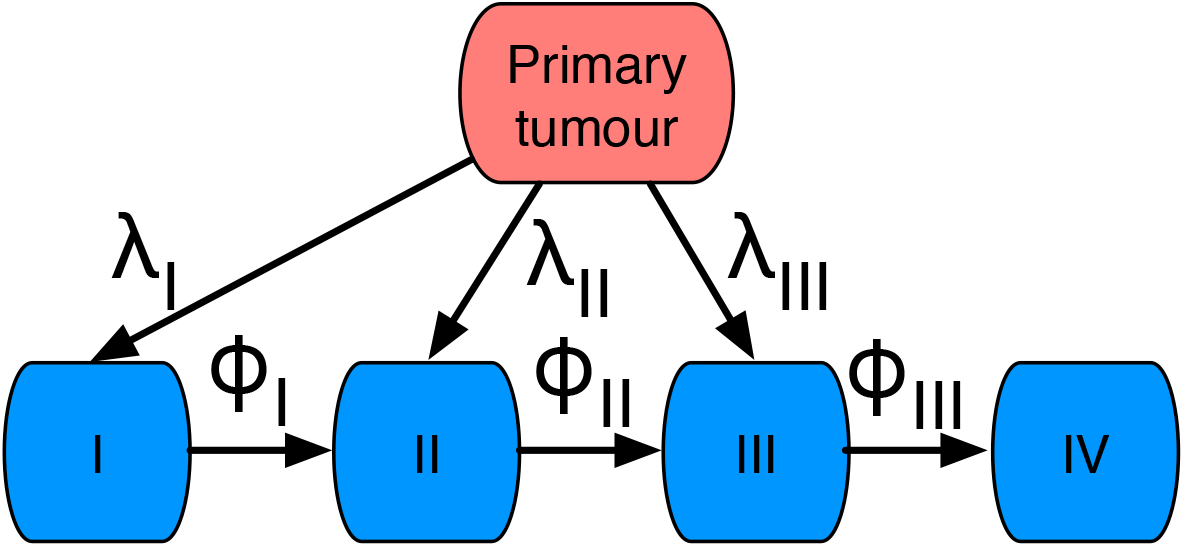
Schematic of the flow of metastatic cells in the case of tongue cancer. The cancer cells flow from the primary tumour to lymph node station I,II and III with rates λ_*I*_, λ_*II*_ and λ_*III*_ respectively. The flow between lymph node stations is defined by the rates *ϕ_I_*, *ϕ_II_* and *ϕ_III_*.

Given a primary tumour metastases will eventually form in downstream stations, but the dynamics of the model crucially depend on the parameters (λ_*I*_, λ_*II*_, λ_*III*_, *ϕ_I_, ϕ_II_, ϕ_III_*). The aim is therefore to estimate these from clinical data. Our data set contains 141 patients with carcinoma of the oral tongue diagnosed between 2004 and 2014 and treated at the department of Oncology at Sahlgrenska University Hospital in Gothenburg (see Methods and Supplementary Information). For each patient we have information about the stage of the primary tumour and the presence/absence of metastasis in LN station I-IV.

In our model we assume that secondary seeding is responsible for all metastases that occur in sites not directly connected to the primary tumour. This implies that we expect all patients with station IV positive to also be positive for station III. This is true for all but 2% of the cases (3 patients), and we therefore exclude these cases from further analysis. Another option would be correct for potentially undetected micrometastases in station III, but we decided for the more cautious option of exclusion.

Given the assumption outlined above about a constant downstream flow of cancer cells and the necessity of secondary seeding the state of the system evolves according to a continuous-time Markov chain with state space that corresponds to all possible states of metastatic spread (see Methods for details). The flow rates in the network dictate transition rates for the Markov chain and the probability distribution over the states changes according to a master equation. Since we know the initial state of the system (no metastasis at tumour initiation) we can, given a set of parameter values, numerically solve the master equation and obtain the probabilities of all the metastatic states for all future times. This allows us to estimate the model parameters by comparing the metastatic state of each patient with solutions to the master equation and computing the likelihood of the data given certain parameter values (see Methods).

In order to visually compare the parametrised model outcome to the clinical data we transform the data in the following way. Let us first focus on station I and let 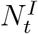 denote the number of patients with stage t primary tumours that are positive in station I. Here *t* can take the values 1, 2, 3,4 corresponding to primary tumour stage T1, T2, T3 and T4. Let *M_t_* denote the total number of patients with stage *t* disease. The fraction of patients with stage *t* tumour with positive lymph node at station I is then given by

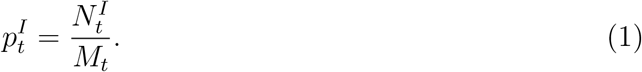

The same procedure is applied to stations II-IV yielding stage-dependent fractions.

The fraction 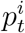 can be interpreted as the probability of finding a patient with stage *t* disease with a metastasis in LN station *i*. This quantity can readily be calculated from the model (see Methods for details).

A comparison of the clinical data and the model fit is shown in figure 2, where each panel correspond to a LN station (I-IV). The estimated dissemination rates are shown in table I together with 95 % confidence intervals obtained using parametric bootstrap (see Methods). It is worth noting that the confidence intervals for *ϕ_I_* and *ϕ_III_* obtained from bootstrapping shows that the variability in flow from the station I to station II is large, ranging from practically zero to 0.42, the largest of all rates. Also the flow from station III to station IV exhibits a large variability. With the current amount of the data we can conclude that flow from station II to III has a large impact on the metastatic process, whereas we are unable to ascertain the importance of flow from station I to II and from III to IV.

**FIG. 2.**
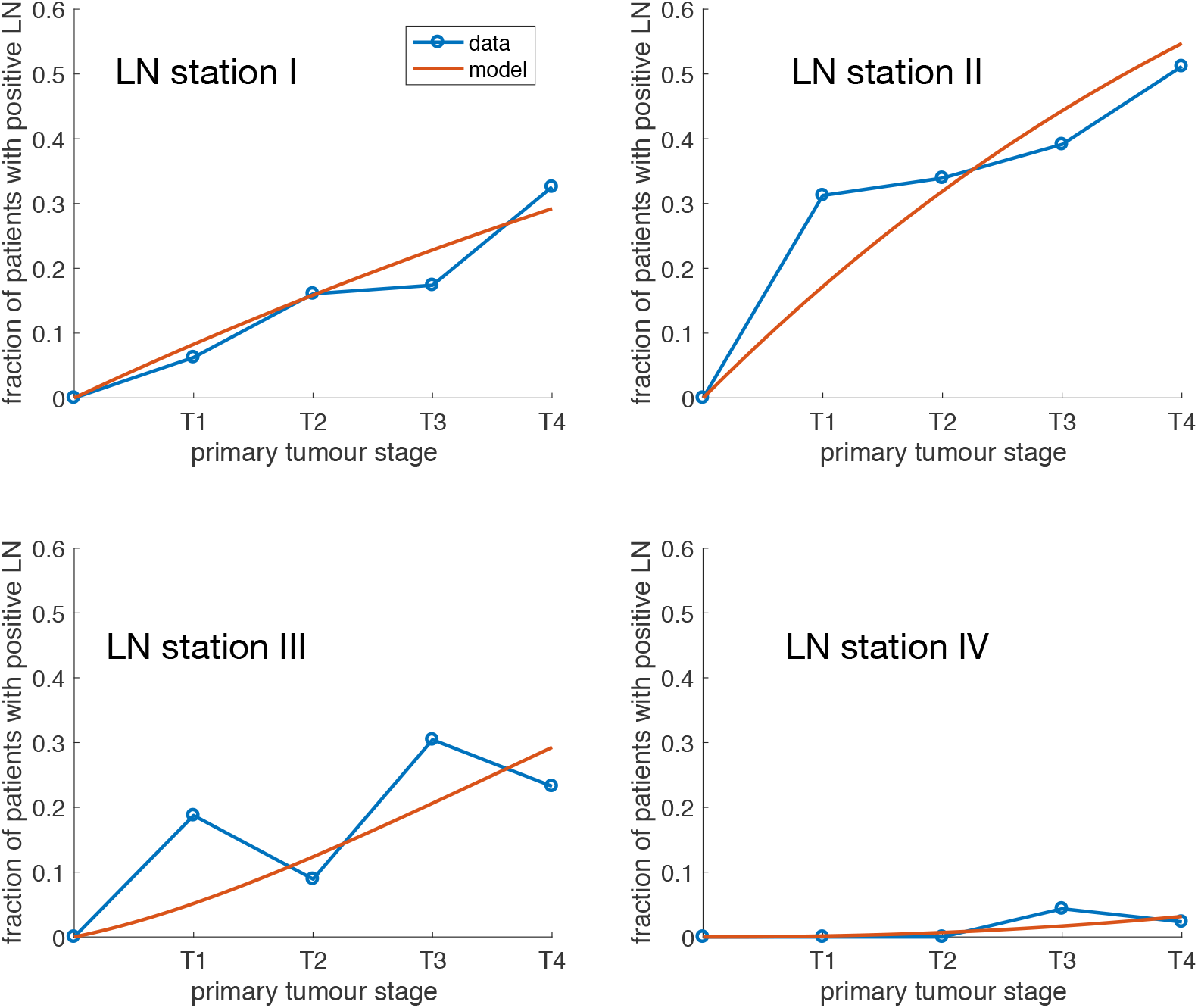
The data and model predictions for metastatic spread to lymph node station I-IV for primary tongue cancer. Each panel corresponds to a lymph node station and shows the probability of finding a metastases in the lymph node as a function of the primary tumour stage. The parameters of the model are given in table I.

**TABLE I.** The parameter estimates for lymphatic spread of tongue cancer.

| Parameter | estimated value | 95 % CI |
| --- | --- | --- |
| $\lambda_I$ | 0.09 | [0.06, 0.12] |
| $\lambda_{II}$ | 0.19 | [0.14, 0.23] |
| $\lambda_{III}$ | 0.04 | [0.02, 0.06] |
| $\phi_I$ | 0.09 | $[9.7 \times 10^{-11}, 0.42]$ |
| $\phi_{II}$ | 0.21 | [0.10, 0.33] |
| $\phi_{III}$ | 0.07 | $[7.5 \times 10^{-10}, 0.15]$ |

### Metastasis in ovarian cancer

Ovarian cancer has the highest mortality rate of the gynecological cancers and a majority of patients are diagnosed in an advanced stage [42]. Ovarian cancer predominantly metastasises within the peritoneal cavity and through the pelvic lymph nodes [43]. In the peritoneal cavity cancer cells metastasize through a process commonly described as transcoelomic dissemination, where the cancer cells loose cell-cell contact and exfoliate into the peritoneal cavity. They float in the peritoneal fluid and are spread across the peritoneal cavity, where they attach to the peritoneal organs and form a metastatic tumour [44]. Ascites produced in the peritoneal cavity is drained through lymph vessels in the diaphragm [45], enabling cancer cells to enter into the blood circulation. Historically, hematogenous metastasis has been regarded as occurring only in late stages of ovarian cancer. Recent work however, suggest that this mode of dissemination may be more common than previously thought [46]. In the case of distant metastases, the most common sites are liver, pleura, lung, central nervous system and skin [47].

The data set on ovarian cancer was obtained from the SEER-database (see Methods). As of 2010 SEER contains information about the presence or absence of metastases at diagnosis in liver, lung, brain and bone. The status of regional LNs (including the pelvis and diaphragm) is also available.

Given the available data we did not try to model transcoelomic dissemination, and instead focused on dissemination to local LNs and hematogenic spread to the organs represented in the SEER database. From a primary tumour located in the ovaries cancer cells can thus either spread to regional LNs or via the venous blood vessels to the lungs (see fig. 3). Metastases in regional LNs also allow for dissemination to the lungs, and from there metastatic lesions shed CTCs to all organs of the body, including the liver, bone and brain (for which we have data).

**FIG. 3.**
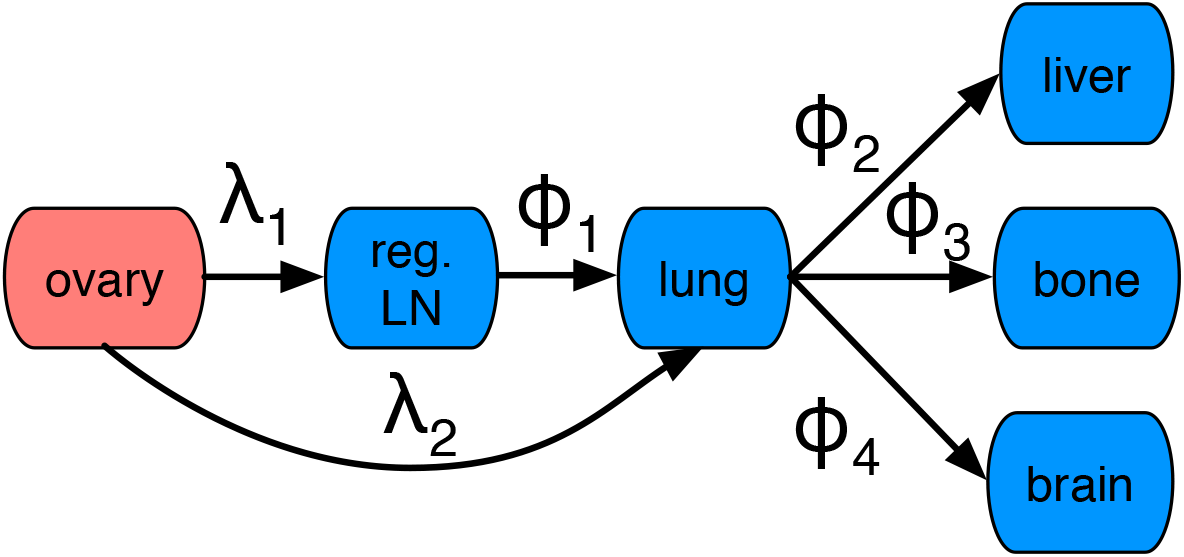
The dissemination network for ovarian cancer. The tumour cells spread either via regional lymph nodes or directly to the lung where they form metastases. From there further dissemination to the liver, bone and brain occurs.

Again our aim is to estimate the dissemination parameters (λ_1_, λ_2_, *ϕ*_1_, *ϕ*_2_, *ϕ*_3_, *ϕ*_4_) from clinical data. In this case the data set is considerably larger containing 16 055 patients diagnosed with ovarian cancer. Again we exclude patients that exhibit skip metastases, leaving us with 15 536 cases (3% of patients are excluded).

The primary tumour stage for this data is more refined and each T-stage is divided into three substages a,b and c, giving us in total 9 different stages (T1a-T3c). In order to fit the parameters we apply a similar maximum likelihood methods as for the previous data set (see Methods).

We estimate the parameters of this model in precisely the same way as for the tongue cancer model (see Methods). Again we compare the parametrised model to the clinical data by calculating the probability of finding a metastasis in each of the sites considered (regional LN, lung, liver, brain and bone) as a function of the primary tumour stage, using equation (1). A comparison between the data and the model is shown in fig. 4, which shows that the model is able to recapitulate the overall behaviour of the data. Due to the low incidence for bone and brain (in total 22 and 5 cases respectively) the data is quite noisy and the model represents a poor fit for those sites. The parameter values are collected in table II together with 95 % confidence intervals obtained using parametric bootstrap (see Methods). For this data set the uncertainty in the inferred paramter values is considerably smaller, which can be attributed to the almost hundred-fold larger data set.

**FIG. 4.**
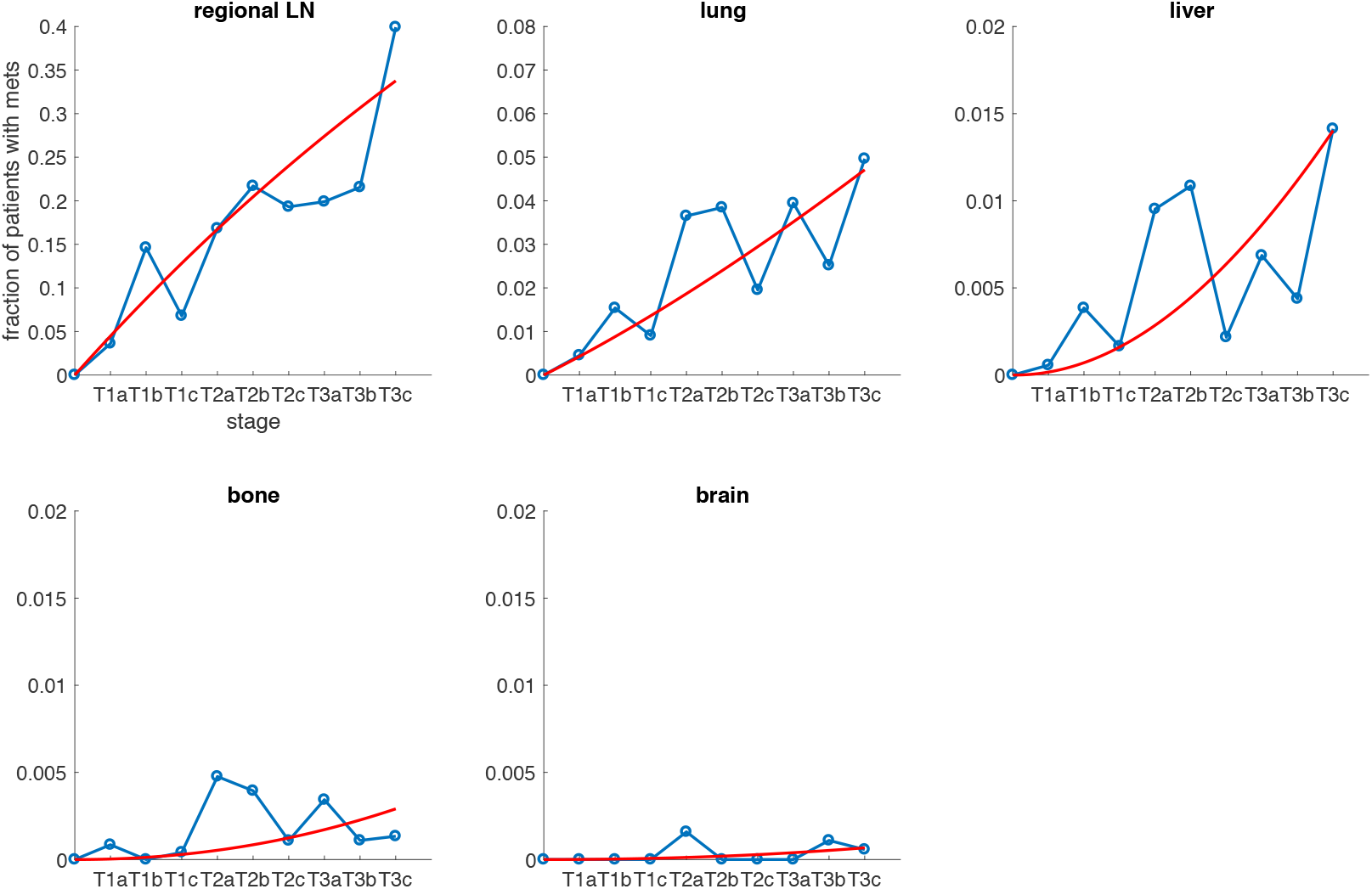
The data and model predictions for metastatic spread to regional lymph nodes and distant sites for primary ovarian cancer. Each panel corresponds to a site/organ and shows the probability of finding a metastases at the site as a function of the primary tumour stage. The parameters of the model are given in table II.

**TABLE II.** The parameter estimates for the metastatic dissemination of ovarian cancer.

| Parameter | estimated value | 95 % CI |
| --- | --- | --- |
| $\lambda_1$ | 0.046 | [0.0446, 0.0476] |
| $\lambda_2$ | 0.004 | [0.0036, 0.0046] |
| $\phi_1$ | 0.007 | [0.0054, 0.0094] |
| $\phi_2$ | 0.092 | [0.0796, 0.1071] |
| $\phi_3$ | 0.015 | [0.0106, 0.0216] |
| $\phi_4$ | 0.004 | [0.0008, 0.0062] |

## DISCUSSION

We have presented a novel method for inferring the rates of metastatic dissemination, and shown that one can obtain reliable estimates (with small CIs) for both lymphatic and hematogeneous spread. Our work builds on previous mathematical models [34], but with additional biological knowledge we simplify the model structure and make parameter estimates more reliable. Firstly, we make use of the fact that high filtration rates imply that secondary seeding is responsible for metastatic spread beyond the first capillary bed/lymph node. This implies that we do not need to consider all possible links between the sites (e.g. no direct link between ovary and liver). Secondly, we make use of known anatomical structures and flow directions to further prune the network (e.g. the flow in the lymphatic system dictates the topology of the tongue cancer network, see fig. 1). Lastly, we make use of primary tumour stage as a proxy for time, which means that we can resolve the data temporally. This implies that we do not have to rely on assumptions about stationarity of the underlying metastatic process, and instead fit the parameters for the time-dependent problem. We assumed that tumour stage maps linearly to an arbitrary time scale, which implies that the inferred dissemination rates are estimated with arbitrary and unknown units. In order to investigate how this assumption affected the inferred parameter values we also considered a model with exponential growth of the primary tumour (see Methods). That model resulted in similar parameter values, which suggest that our approach is robust to assumptions about tumour growth dynamics.

Our method makes it possible to infer dissemination rates between sites and also their confidence intervals. The model does not fit the data perfectly, and although this may have to do with the scarcity of the data set, we cannot exclude that this may be due to some limitations of the assumptions used in the model. Traditionally this type of data is analysed by looking at incidence rates [48]. Our analysis goes beyond this by disentangling incidence rates into dissemination rates between different lymph node stations/organs. This means that we can quantify processes that at a first glance seem inaccessible, but which appropriate assumptions and modelling techniques can help us reveal.

It could be argued that we have used an excessively complex model to fit a data set, which could be described with four (and five) straight lines (see fig. 2 and fig. 4), and hence four (and five) parameters corresponding to the slopes of the lines. However, such a model would have no connection to the underlying biology and would be unable to say anything about the dynamics that generated the data. In our model on the other hand the parameters have an immediate biological interpretation. They correspond to the (relative) flow rates of cancer cells between different anatomical sites, numbers that could be of interest to clinicians when deciding on the extent of surgical resection or radiotherapy. The magnitudes of the flow rates also reveal the importance of different routes of spread. For example for ovarian cancer we can conclude that the rate of metastasis formation in the regional lymph nodes is ten times higher than the direct spread to the lungs.

The flow rates in our model factor in not only physical flow, but also the ability of cancer cells to survive during transport to the target site, and their ability to form metastases in the target site. Assuming that the survival in the circulatory system is independent of target site, knowledge of physical flow would make it possible to estimate the relative rate of metastasis formation in different target sites. This would correspond to effect of the “soil” in the well-established seed-soil hypothesis [49, 50]. Unfortunately, lymphatic flow is difficult to measure [51], and there are currently no estimates of lymphatic flow in the head and neck region. Values of relative blood flow are however readily available [52], and we can use these to calculate soil-effect for ovary cancer with respect to liver, bone and brain. The relative blood flow to these organs is given by 6.5%,a 5% and 1.2% and by dividing these flow rates with *ϕ*_2_, *ϕ*_3_ and *ϕ*_4_, we get relative metastasis formation rates of 1.42, 0.3 and 0.33. From this we can conclude that the rate of metastasis formation in the liver is roughly five fold higher compared to bone and brain. This suggests that ovarian cancer cells are considerably better at colonising the liver compared to bone and brain, which is in agreement with previous data [53].

We did encounter a couple of issues when it comes to parameter identifiability. In the case of tongue cancer our model was unable to accurately identify the rates *ϕ_I_* and *ϕ_III_*. In the case of *ϕ_I_* the problem arises because we are dealing with a site (station II) which has flow from both the primary tumour and an upstream site (i.e. station I). With current size of the dataset (*n* = 141 patients) we are unable to obtain accurate estimates of the flow into this station. With a larger dataset we would most likely find more patients which are only positive for station II making it possible to obtain better estimates for λ_*II*_ and consequently *ϕ_I_*. The large confidence interval obtained for *ϕ_III_* is due to the low number of cases with metastases in station IV. In the bootstrap procedure we generate synthetic data based on the estimated parameter values, and since the actual number of patients with station IV positive is small (only two cases) we sometimes generate data without any prevalence of metastasis in station IV. This results in estimating the flow rate to *ϕ_III_* = 0, which in turn leads to a large confidence interval. This problem does not appear for the ovarian model since, although the inferred parameter values are small, the large number of patients in the data set imply that we almost surely generate synthetic data with some patients positive for metastases in the brain or bone.

From a clinical point of view, this work could be of importance, by contributing to an increased possibility to predict the risk of future regional and/or distant metastases. Especially so in the current era, with new treatment modalities emerging and a current development towards more individualised treatment programs. Although the models are parametrised with population level data they might still be used in order to make predictions on the individual level. For example if a patient is diagnosed with tongue cancer of a specific stage the parametrised model could provide probabilities of different metastatic states, which could be factored in with other clinical data to guide treatment. It would also be possible to include the effect of treatment in the model (e.g. radiation), which would reveal potential benefits of treating metastatic sites and making the model better suited to describe clinical procedure.

In conclusion we believe that this framework for analysing metastatic spread, which incorporates known anatomical constraints and a temporal dimension, allows for novel insights and will hopefully be of assistance to both cancer biologists and clinicians in the future.

## METHODS

### Ethics Statement

For the tongue cancer data approval has been granted by Regional ethical review board in Gothenburg.

### Data

#### Tongue cancer

After approval from the Regional ethical review board in Gothenburg, data was obtained from the register of the department of Oncology at Sahlgrenska University Hospital in Gothenburg. Data for all patients diagnosed with carcinoma of the oral tongue between 2004 and 2014 were extracted, a total number of 141 cases. Information about primary tumour, LN metastases and distant metastes was registered according to the TNM classification system (AJCC 6th edition until 2009 and 7th edition from 2010 on). T-stage varied from T1 to T4, 78 patients had regional LN metastes (N1 - N3) and only two patients had distant metastasis (M1), in the lung. For patients with positive nodal status, we collected information about in which lymph node levels (I-V) metastases were present.

Only one patient exhibited metastasis in lymph node level V, and therefore this level was excluded from the analysis. Information about involved LN levels was in approximately 50 % of the cases retrieved from radiological examination (CT scans or magnetic resonance imaging), why patients received primary radiotherapy or surgery was performed without neck dissection. For the remaining part this information was obtained from the pathology reports performed after primary surgery including supraomohyoid neck dissection. For each patient we had the following information:

- T-stage (derived AJCC 6th and 7th edition)
- Presence/Absence of regional LN metastases (calculated from N-stage)
- Localization of regional LN metastases (lymph node levels I-VI)

For further information about the data set please see Supplementary Material.

#### Ovarian cancer

The data was obtained from the SEER*Stat case listing database *Incidence – SEER 18 Regs Research Data + Hurricane Katrina Impacted Louisiana Cases, Nov 2016 Sub (2000-2014)*, which is available at https://seer.cancer.gov/seerstat/. We extracted all cases of ovarian cancer from 2010-2014 with recorded metastases in liver, lung, brain and bone. From this set of patients we selected those with a T-stage (derived AJCC 7th edition) which was in the range T1a to T3c. Information about the presence or absence of regional lymph node metastases was obtained by looking at the N-stage (derived AJCC 7th edition). We considered patients with N0 to be free of regional LN metastases while all other stages to be indicative of positive LN. The total number of cases that met the above critera was 16 055 and for each case we had the following information:

- T-stage (derived AJCC 7th edition)
- Presence/Absence of regional LN metastases (calculated from N-stage)
- Presence/Absence of metastases in lung (CS-mets at DX-lung)
- Presence/Absence of metastases in liver (CS-mets at DX-liver)
- Presence/Absence of metastases in bone (CS-mets at DX-bone)
- Presence/Absence of metastases in brain (CS-mets at DX-brain)

### Mathematical model

The model consists of *N* nodes each representing a specific site/organ (excluding the primary site). A node takes the value 0 at time *t* if the site is void of metastases and 1 if the site contains one or more metastases. We assume that once a site has become positive it will remain so for all future times. Since we have *N* nodes and each can be in two states we have a state space for the entire network that contains 2^*N*^ states. Each state is denoted by a binary string (e.g. 0100 correspond to site 1,3 and 4 being negative and site 2 being positive), but for notational simplicity we enumerate them with integers from 1 to 2^*N*^.

The flow rates in the anatomical network dictate transition rates between states. We assume that a site becomes positive at a rate which equals the sum of flow rates from all positive upstream sites. Since the transition rates only depend on the current state (and not the history) the system can be described as a continuous-time Markov chain with state space {0,1}^*N*^. If we let *P_i_(t)*, where *i* = 1,…, 2^*N*^, denote the probability of being in state *i* at time *t*, then these probabilities evolve according to the master equation [54]:

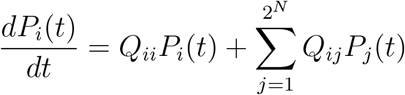

where *Q_ii_* ≤ 0 is the rate of leaving state *i* and *Q_ij_* is the rate of moving from state *j* to *i*. Since we know that the patient is free of metastasis at tumour intitiation (*t* = 0), we know that the initial condition for the master equation is given by *P*_1_(*t*) = 1 and *P*_*i*_(*t*) = 0 for 1 < *i* ≤ 2^*N*^. Numerical solutions of this equation will be used in order to parametrise the two models.

Since we only have access to the primary tumour stage of the patients in the two data sets we make the following assumption about the mapping from tumour stage at diagnosis to time from tumour intitiation to diagnosis. For simplicity we assume that the tumour radius grows linearly with time, which is a simplification, since it is known that volume and hence radius grows non-linearily with time [37].

Tumour stage is informed by the linear size of the lesion [55], e.g. for tongue cancer T1 corresponds to a primary tumour less than 1 cm in diameter and T2 is a tumour larger than 2 cm, but less than 4 cm. This means that there is a strong correlation between tumour stage and thickness [56]. We therefore assume that time from tumour initiation depends linearly on the stage. Since we have no exact information about the details of this mapping we will for simplicity consider an arbitrary time scale and simply let the stage correspond directly to time since initiation. Below we challenge this assumption by considering a model in which the primary tumour volume grows exponentially.

#### Tongue cancer

We let lymph node station I-IV correspond to node 1-4 in the network. Since we have four nodes we will have a state space of 2^4^ = 16 metastatic states, but 4 of these will be inaccessible due to topology of the anatomical network (e.g. 1001 is not attainable) and we are left with 12 states. The transition rates are shown in fig. 5, where the nodes now represent states and the arrows show possible transitions and their corresponding ates.

**FIG. 5.**
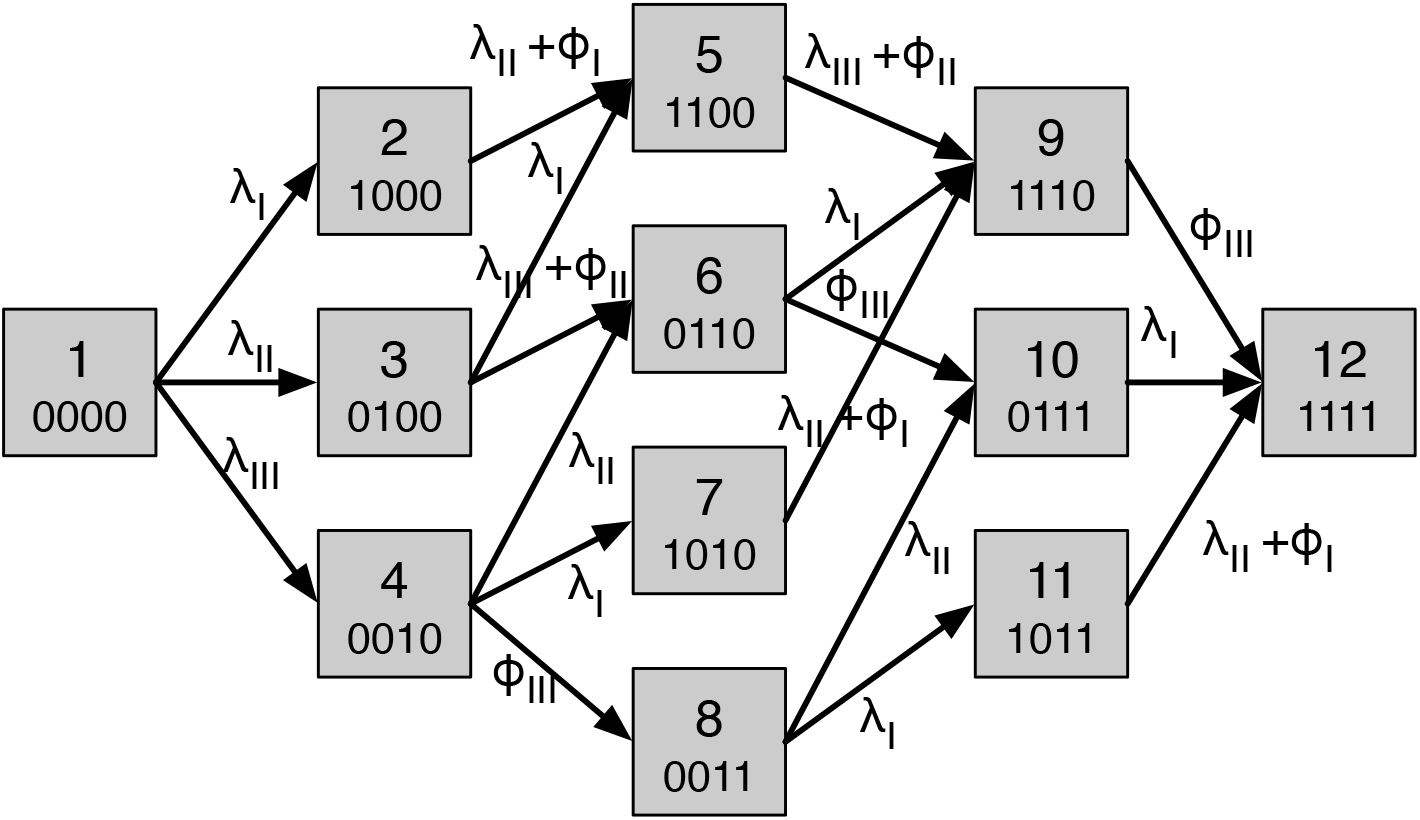
The state transition diagram for the tongue cancer model. Each node corresponds to a possible metastatic state and is represented by a binary string which signifies which lymph node station is positive or negative. For convenience the nodes are named from 1 to 12. The rates are obtained from the flow rates in the anatomic network 1 by considering the rate at which individual sites become positive based on the state of the other sites.

The transition matrix is given by

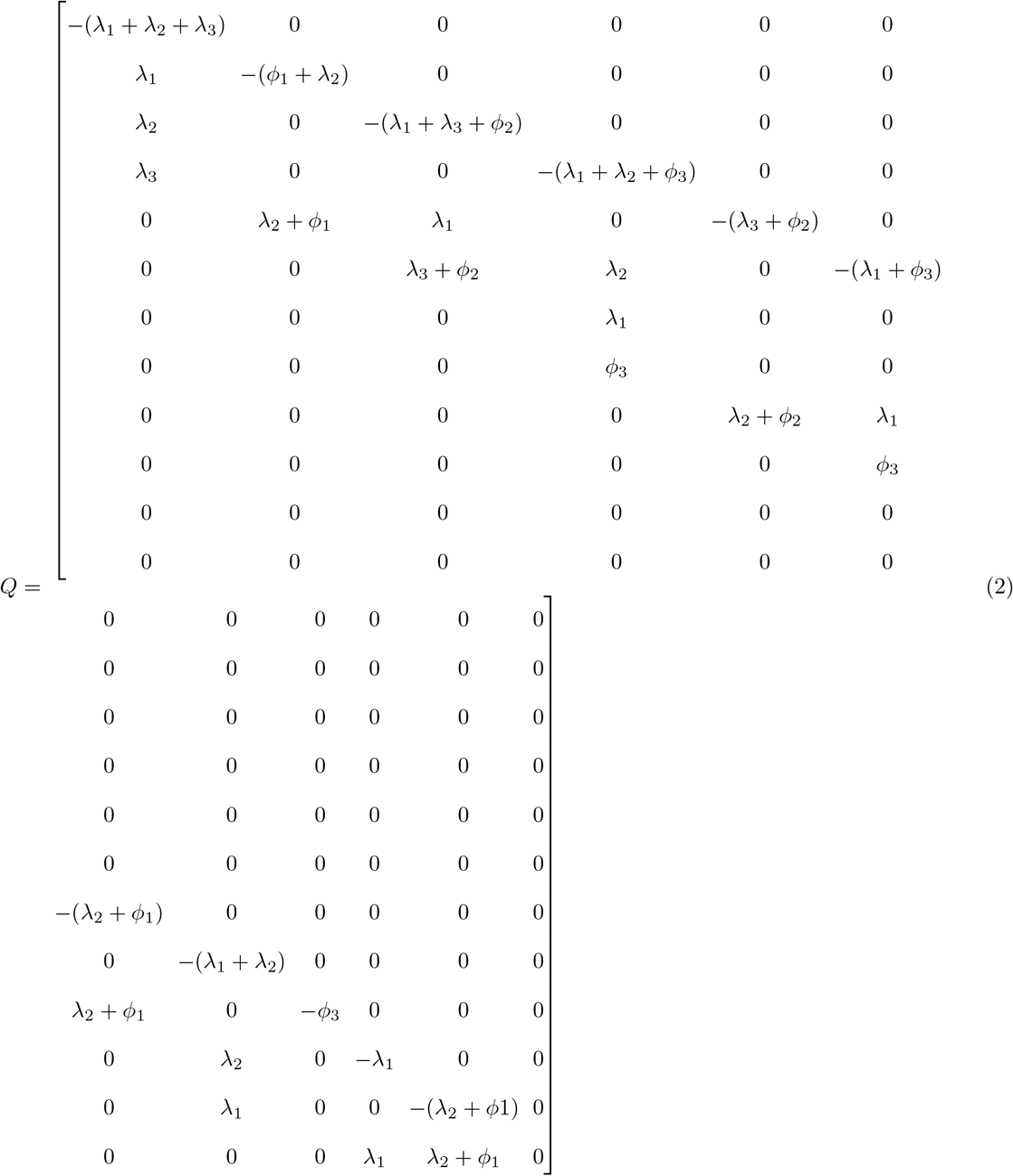

With **P** = [*P*_1_, *P*_2_…, *P*_12_]^*T*^ we can write the master equation as

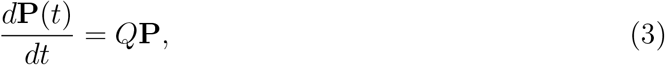

which we solve numerically using a first-order Euler forward scheme with initial condition *P*_1_(*t*) = 1 and *P_i_(t)* = 0 for 1 < *i* ≤ 12, which corresponds to no metastasis at tumour initiation.

To estimate the parameters we make use of a maximum likelihood method. Preferably we would like to have the time until onset of metastasis for each station in each patient, but we only have access to data obtained at diagnosis. This information can be used in the following way: Assume that we observe a patient with a stage *τ* tumour that has a positive lymph node at station I and all other stations negative, i.e. the patient is in state 1000. The probability of this occurring is according to the model given by *P*_2_(*τ*). Now the value of *P*_2_(*τ*) will depend on the parameters of the model and we therefore write *P*_2_(*τ, θ*), where *θ* = [λ_*I*_, λ_*II*_, λ_*III*_, *ϕ_I_, ϕ_II_, ϕ_III_*]. The likelihood of the entire clinical data set can now be written

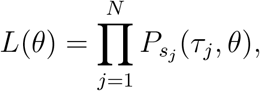

where *s_j_* is the metastatic state of patient *j*, which can take values 1,2,…,12, and *τ_j_* ∈ {1, 2, 3, 4} is the primary tumour stage of that patient. In order to find the parameters *θ* that best fit the data we minimise the negative likelihood — *L*(*θ*) using the MATLAB function `fminsearch` applied to a numerical solution of (3). This gives us a maximum likelihood estimate 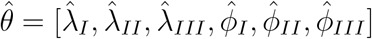.

A solution to eq. (3) with parameter values obtained from the maximum likelihood method is shown in fig. 6. We note that the probability of the metastasis free state (0000) decays over time while all other probabilities increase over the considered time frame. For large (and unrealistic) times all probabilities will tend to zero expect the state corresponding to metastases in all sites (1111), which will tend to unity.

**FIG. 6.**
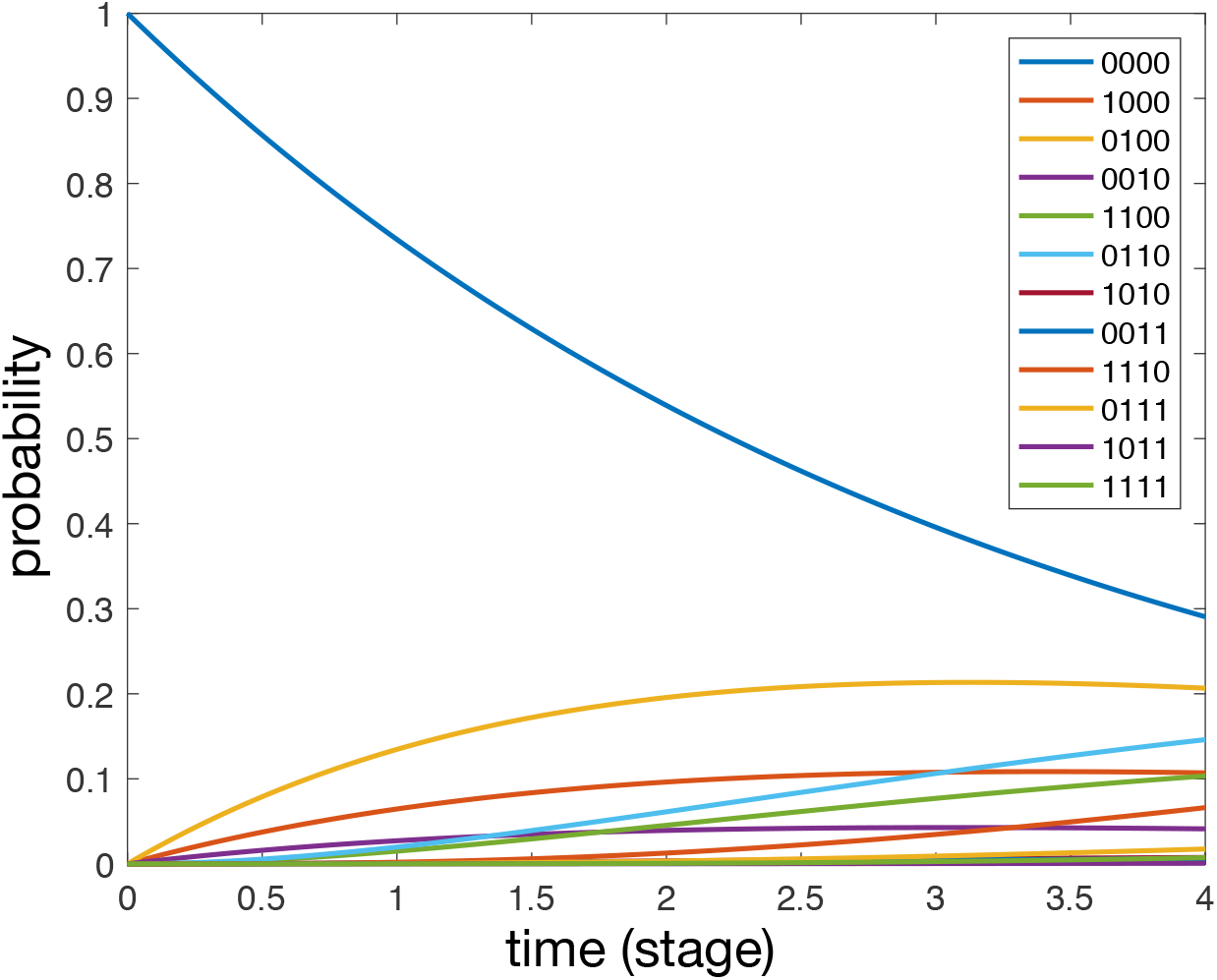
A solution to eq. (3) with parameter values obtained from the maximum likelihood method. Each curve correspond to specific state of the system.

Confidence intervals for 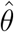 are calculated using parametric bootstrapping with 100 samples [57]. This is a numerical method in which the parameter estimates obtained from the maximum likelihood method are used for generating synthetic data, from which new maximum likelihood estimates are obtained. Repeating this process results in an empirical distribution for each parameter, and the confidence intervals are calculated as the 5% and 95% quantiles of the empirical distributions.

The probability that station I is positive (the solid curve in fig. 2 top-left panel) can be calculated from the model by summing over the probability of all metastatic states which have 1 in the first position of the binary string, i.e.

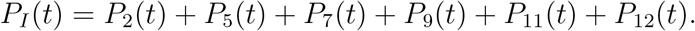

In the same way we can write down

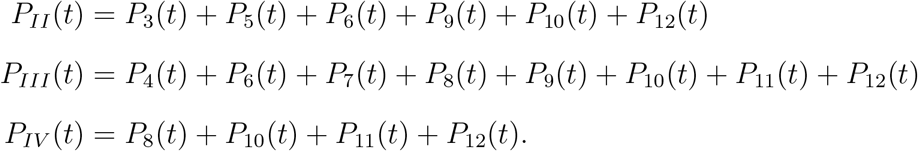

#### Ovarian cancer

We let regional lymph nodes correspond to node 1 in the network, lung to node 2, liver to node 3, bone to node 4 and brain to node 5. Since we have five nodes we will have a state space of 2^5^ = 32 metastatic states, but 14 of these are inaccessible due to topology of the anatomical network and we are left with 18 states. The transition rates are shown in fig. 7, where the nodes now represent states and the arrows show possible transitions and their corresponding rates.

**FIG. 7.**
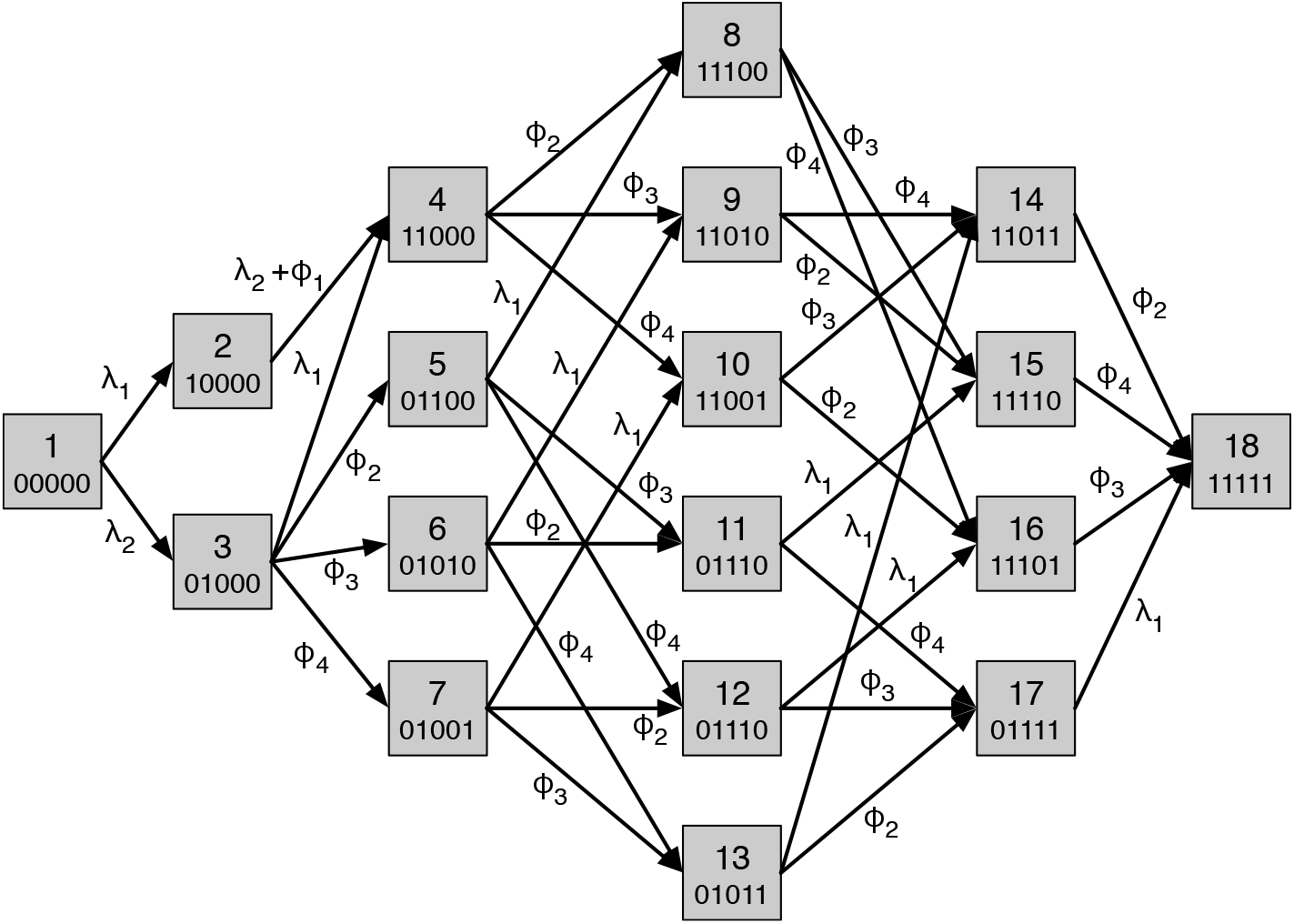
The state transition diagram for the ovarian cancer model. Each node corresponds to a possible metastatic state and is represented by a binary string which signifies which organ is positive or negative for metastases. For convenience the nodes are named from 1 to 18. The rates are obtained from the flow rates in the anatomic network (fig. 3) by considering the rate at which individual sites become positive based on the state of the other sites.

We now have an 18-dimensional probability vector **P** = [*P*_1_, *P*_2_…. *P*_18_]^*T*^ and we can again write down the master equation

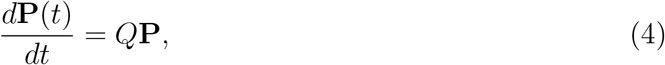

where the matrix *Q* is defined as above and can be obtained from the state transition diagram fig. 7. We proceed as above and formulate the maximum likelihood problem that we solve numerically using MATLAB.

A solution to eq. (4) with parameter values obtained from the maximum likelihood method is shown in fig. 8. We note that the dynamics are dominated by the metastasis free state (00000) and the state with only regional LNs positive (01000), while all other states have probabilities less than 0.02. For large (and unrealistic) times all probabilities will tend to zero expect the state corresponding to metastases in all sites (11111), which will tend to unity.

**FIG. 8.**
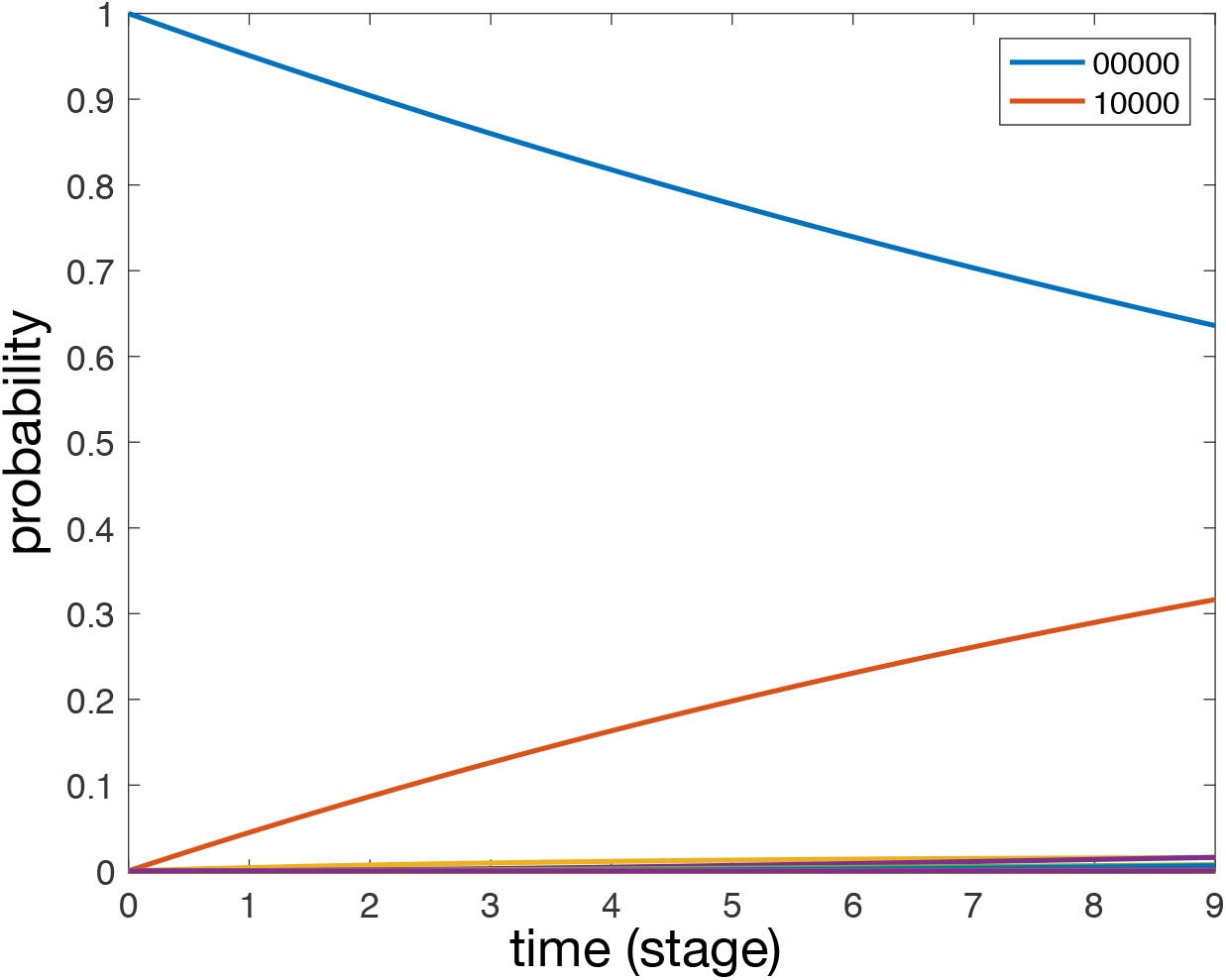
A solution to eq. (4) with parameter values obtained from the maximum likelihood method. Each curve correspond to specific state of the system.

The probability of finding a patient with metastasis in the regional lymph nodes can be expressed by summing the probability over all states in which node 1 is positive

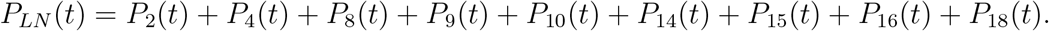

In the same way we can write down

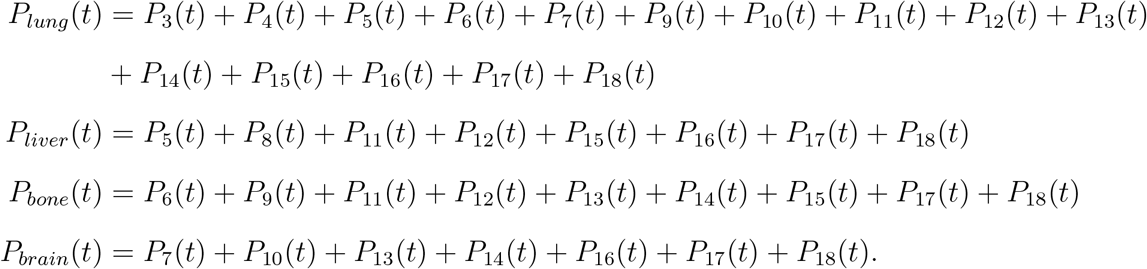

#### Exponential tumour growth

In order to assess the impact of our assumption that primary tumour stage can be used as a proxy for time we also consider a model of tongue cancer metastasis in which the times corresponding to the different tumour stages are calculated assuming an exponential growth in tumour volume.

If we assume that tumour volume grows according to

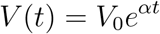

Assuming the tumour to be spherical, the radius, which is used for tumour staging [55], grows as

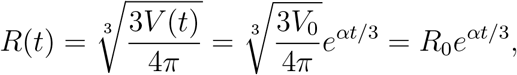

with 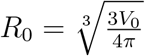. If we let *R_i_* denote the radius of a stage *i* tumour we get that the time at which stage *i* is reached is given by

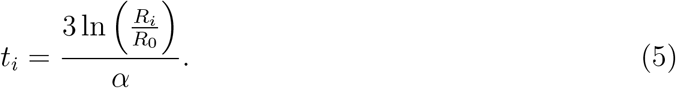

The initial radius of the tumour is set to be equal to the volume of a single cell *R*_0_ = 25 μm and the growth rate is estimated from clinical studies to 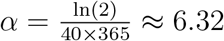 year^−1^ [58].

The clinical guidelines for determining tumour stage of tongue cancer involve the size of the lesion but also other factors such as degree of invasion of surrounding tissue [55]. We simplify this by considering the following mapping from tumour stage to tumour size. We let a T1 tumour have a radius of 2 cm, a T2 tumour have radius 4 cm, a T3 tumour have radius 6 cm and a T4 tumour have radius 8 cm. Of course this is highly simplistic but at least provides us with a reasonable quantification.

This leads us to a mapping from tumour stage to time from initiation which is shown in table III.

**TABLE III.** The mapping from tumour stage to time since initiation for tongue cancer assuming exponential growth.

| tumour stage | time since initiation (years) |
| --- | --- |
| 1 | 3.17 |
| 2 | 3.50 |
| 3 | 3.69 |
| 4 | 3.83 |

We now use these values (instead of the identity mapping from stage to time) in our maximum likelihood estimation and obtain the parameter values shown in table IV. The parameters obtained with the exponential growth model are generally slightly smaller, although *ϕ_I_* is an exception with a larger value compared to the linear model. But more importantly the order of their magnitudes is the same for both models, i.e. *ϕ_II_* > λ_*II*_ > *ϕ_I_* > λ_I_ > *ϕ_III_* > λ_*III*_ for both models.

**TABLE IV.** The parameter estimates for lymphatic spread of tongue cancer.

| Parameter | linear model | exponential model | ratio |
| --- | --- | --- | --- |
| $\lambda_I$ | 0.09 | 0.06 | 0.73 |
| $\lambda_{II}$ | 0.19 | 0.13 | 0.71 |
| $\lambda_{III}$ | 0.04 | 0.03 | 0.70 |
| $\phi_I$ | 0.09 | 0.11 | 1.19 |
| $\phi_{II}$ | 0.21 | 0.17 | 0.82 |
| $\phi_{III}$ | 0.07 | 0.06 | 0.82 |

This suggests that our model of metastatic spread is robust different assumption about tumour growth dynamics.

## References

[1] K. A. Price and E. E. Cohen, Current treatment options in oncology 13, 35 (2012).

[2] K. Deng, C. Yang, Q. Tan, W. Song, M. Lu, W. Zhao, G. Lou, Z. Li, K. Li, and Y. Hou, Gynecologic oncology 150, 460 (2018).

[3] I. O. Bello, Y. Soini, and T. Salo, Oral oncology 46, 636 (2010).

[4] P. Dahm-Kähler, C. Borgfeldt, E. Holmberg, C. Staf, H. Falconer, M. Bjurberg, P. Kjälhede, P. Rosenberg, K. Stålberg, T. Högberg, et al., Gynecologic oncology 144, 167 (2017).

[5] A. Du Bois, A. Reuss, E. Pujade-Lauraine, P. Harter, I. Ray-Coquard, and J. Pfisterer, Cancer 115, 1234 (2009).

[6] G. Curigliano, C. Criscitiello, A. Esposito, and G. Pruneri, The Breast 31, 303 (2017).

[7] D. M. Trifiletti, A. Smith, N. Mitra, S. Grover, J. N. Lukens, R. B. Cohen, P. Read, W. M. Mendenhall, A. Lin, and S. Swisher-McClure, Journal of Clinical Oncology 35, 1550 (2017).

[8] L. Weiss, K. Haydock, J. Pickren, and W. Lane, The American journal of pathology 101, 101 (1980).

[9] S. K. Mukherji, D. Armao, and V. M. Joshi, Head & Neck 23, 995 (2001).

[10] D. Hanahan and R. A. Weinberg, cell 144, 646 (2011).

[11] R. Govindan and V. T. DeVita, DeVita, Hellman, and Rosenberg’s Cancer: Principles & Practice of Oncology Review (Lippincott Williams & Wilkins, 2009).

[12] G. P. Gupta and J. Massagué, Cell 127, 679 (2006).

[13] S. Valastyan and R. A. Weinberg, Cell 147, 275 (2011).

[14] T. P. Butler and P. M. Gullino, Cancer research 35, 512 (1975).

[15] M. Al-Hajj, M. S. Wicha, A. Benito-Hernandez, S. J. Morrison, and M. F. Clarke, Proceedings of the National Academy of Sciences 100, 3983 (2003).

[16] Y. Okumura, F. Tanaka, K. Yoneda, M. Hashimoto, T. Takuwa, N. Kondo, and S. Hasegawa, The Annals of thoracic surgery 87, 1669 (2009).

[17] L. Weiss, Clinical and Experimental Metastasis 10, 191 (1992).

[18] J. G. Scott, A. G. Fletcher, P. K. Maini, A. R. Anderson, and P. Gerlee, European journal of cancer 50, 3068 (2014).

[19] Scott, D. Basanta, A. Anderson, and P. Gerlee, Journal of The Royal Society Interface 10, 1 (2013).

[20] S. Meng, D. Tripathy, E. P. Frenkel, S. Shete, E. Z. Naftalis, J. F. Huth, P. D. Beitsch, M. Leitch, S. Hoover, D. Euhus, et al., Clinical cancer research 10, 8152 (2004).

[21] I. Bross, E. Viadana, and J. Pickren, Journal of chronic diseases 28, 149 (1975).

[22] K. P. McMullen, J. J. Chalmers, J. C. Lang, P. Kumar, and K. R. Jatana, World journal of otorhinolaryngology-head and neck surgery 2, 109 (2016).

[23] L. Liotta, G. Saidel, and J. Kleinerman, Biometrics, 535 (1976).

[24] L. Liotta, C. Delisi, G. Saidel, and J. Kleinerman, Cancer Letters 3, 203 (1977).

[25] G. Saidel, L. Liotta, and J. Kleinerman, Journal Of Theoretical Biology 56, 417 (1976).

[26] K. Iwata, K. Kawasaki, and N. Shigesada, Journal Of Theoretical Biology 203, 177 (2000).

[27] E. Baratchart, S. Benzekry, A. Bikfalvi, T. Colin, L. S. Cooley, R. Pineau, E. J. Ribot, O. Saut, and W. Souleyreau, PLoS Computational Biology 11, e1004626 (2015).

[28] S. Benzekry, A. Tracz, M. Mastri, R. Corbelli, D. Barbolosi, and J. M. L. Ebos, Cancer Research 76, 535 (2016).

[29] L. Hanin, K. Seidel, and D. Stoevesandt, Journal of Mathematical Biology 72, 1633 (2015).

[30] H. Haeno, M. Gonen, M. B. Davis, J. M. Herman, C. A. Iacobuzio-Donahue, and F. Michor, Cell 148, 362 (2012).

[31] R. Demicheli, M. W. Retsky, D. E. Swartzendruber, and G. Bonadonna, Annals of oncology: official journal of the European Society for Medical Oncology 8, 1075 (1997).

[32] L. Norton and J. Massagué, Nature medicine 12, 875 (2006).

[33] J. Scott, P. Kuhn, and A. R. A. Anderson, Nature Reviews Cancer, 1 (2012).

[34] P. K. Newton, J. Mason, K. Bethel, L. A. Bazhenova, J. Nieva, and P. Kuhn, PLoS ONE 7, e34637 (2012).

[35] P. K. Newton, J. Mason, K. Bethel, L. Bazhenova, J. Nieva, L. Norton, and P. Kuhn, Cancer Research 73, 2760 (2013).

[36] G. Disibio and S. W. French, Archives of pathology & laboratory medicine 132, 931 (2008).

[37] P. Gerlee, Cancer research 73, 2407 (2013).

[38] R. M. Byers, R. S. Weber, T. Andrews, D. McGill, R. Kare, and P. Wolf, Head & neck 19, 14 (1997).

[39] C. Klein, Nature Reviews Cancer 9, 302 (2009).

[40] H. Wang, C. Zhang, J. Zhang, L. Kong, H. Zhu, and J. Yu, Oncotarget 8, 26368 (2017).

[41] N. Benson, M. Whipple, and I. J. Kalet, AMIA … Annual Symposium proceedings. AMIA Symposium, 31 (2006).

[42] R. L. Siegel, K. D. Miller, and A. Jemal, CA: a cancer journal for clinicians 65, 5 (2015).

[43] N. Tsuruchi, T. Kamura, N. Tsukamoto, K. Akazawa, T. Saito, T. Kaku, N. To, and H. Nakano, Gynecologic oncology 49, 51 (1993).

[44] A. K. Mitra, in Tumor Metastasis (InTech, 2016).

[45] D. S. Tan, R. Agarwal, and S. B. Kaye, The lancet oncology 7, 925 (2006).

[46] S. Pradeep, S. W. Kim, S. Y. Wu, M. Nishimura, P. Chaluvally-Raghavan, T. Miyake, C. V. Pecot, S.-J. Kim, H. J. Choi, F. Z. Bischoff, et al., Cancer cell 26, 77 (2014).

[47] G. Cormio, C. Rossi, A. Cazzolla, L. Resta, G. Loverro, P. Greco, and L. Selvaggi, International Journal of Gynecological Cancer 13, 125 (2003).

[48] M. Qiu, J. Hu, D. Yang, D. P. Cosgrove, and R. Xu, Oncotarget 6, 38658 (2015).

[49] S. Paget, Cancer metastasis reviews 8, 98 (1989).

[50] I. J. Fidler, Nature Reviews Cancer 3, 453 (2003).

[51] C. Blatter, E. F. J. Meijer, A. S. Nam, D. Jones, B. E. Bouma, T. P. Padera, and B. J. Vakoc, Scientific Reports, 1 (2016).

[52] R. W. Leggett and L. R. Williams, Health physics 69, 187 (1995).

[53] G. Disibio and S. W. French, Archives of pathology & laboratory medicine 132, 931 (2008).

[54] C. Gardiner, Stochastic methods, Vol. 4 (Springer-Verlag Berlin Heidelberg, 2009).

[55] S. B. Edge and C. C. Compton, Annals of surgical oncology 17, 1471 (2010).

[56] S. Zia, S. U. Naqvi, H. Adel, S. O. Adil, M. Hussain, et al., Pakistan journal of medical sciences 33, 353 (2017).

[57] C. Z. Mooney, R. D. Duval, and R. Duvall, Bootstrapping: A nonparametric approach to statistical inference, 94–95 (Sage, 1993).

[58] H. Komatsubara, M. Umeda, T. Minamikawa, Y. Ojima, M. Yanagita, T. Shigeta, Y. Shibuya, S. Yokoo, and T. Komori, Journal of Japan Society for Oral Tumors 17, 232 (2005).

